# A Mechanism for the Rare Fluctuation that Powers Protein Conformational Change

**DOI:** 10.1101/2021.09.27.462043

**Authors:** Shanshan Wu, Ao Ma

## Abstract

Most functional processes of biomolecules are rare events. Key to a rare event is the rare fluctuation that enables the energy activation process, which powers the system across the activation barrier. But the physical nature of this rare fluctuation and how it enables barrier crossing are unknown. With the help of a novel metric, the reaction capacity *p*_*C*_, that rigorously defines the beginning and parameterizes the progress of energy activation, the rare fluctuation was identified as a special phase-space condition that is necessary and sufficient for initiating systematic energy flow from the non-reaction coordinates into the reaction coordinates. The energy activation of a prototype biomolecular isomerization reaction is dominated by kinetic energy transferring into and accumulating in the reaction coordinates, administered by inertial forces alone. The two major reaction coordinates move in precise synergy, with one acting as a gating mechanism on the other. This mechanism is enabled by the structural features of biomolecules and may the cause of their unique functions that are not possible in small molecules.

---

Many important functional processes in molecular systems, such as chemical reaction, protein folding, enzymatic reaction, are rare events—events with time scales much slower than elementary molecular motions ^1-3^. A rare event is rare because it requires activation: between the initial (reactant) and final (product) states that define the transition there is an activation barrier much higher than thermal energy that the system must cross, which requires extra energy. In a rare event, there are two characteristic times, one is the long waiting time a molecule spent in the reactant state, the other is the much shorter transition time, in which a molecule moves from the reactant to the product state ^4^.

Predicting the time scales of rare events, i.e. reaction rates, has been a central topic in chemical physics and biophysics since the beginning of studies on rare events. The standard conceptual and theoretical framework is based on transition state theory and Kramers theory, with Grote-Hynes theory an extension of the latter to the non-Markovian regime ^5-12^. The focus of these theories was to estimate the first passage time (i.e. waiting time), which is the foundation for calculating the reaction rate. In recent years, advances in single molecule experiments have enabled measurements of transition time, which motivated considerable theoretical efforts in determining the transition time distribution for model systems ^4,13-15^. On the other hand, there is a compelling need to understand the mechanism of the transition period dynamics, which is of particular importance for biomolecules.

A fundamental difference between biomolecules and small molecules is that the former is much larger and structurally more complex. The added size and structural complexity endow biomolecules with functions that are not possible for small molecules. Aside from the natural interest in understanding how the added complexity leads to the enriched functionality, there is a practical demand for understanding how to modify functions of biomolecules to better suit human needs (e.g. new or improved catalytic power of enzymes) or avoid undesirable consequences (e.g. drug resistance). Either way requires understanding the detailed mechanism of rare events in biomolecules because there is a plethora of tunable factors in a functional process that coordinate with each other to achieve the function of a biomolecule. To modify a function effectively, we need to know what these factors are and how they work together ^3,16^. Only in this way will we be able to tune the proper factors and achieve intended outcomes. Without reliable understanding of the mechanism, the success rate for blind trial-and-error approach will be too low. Therefore, understanding the physical mechanism of rare events is of crucial importance. The essence of a rare event is its transition period.

In the general physical picture of a rare event, a molecule spends a long time in the reactant basin undergoing equilibrium fluctuations and waits for a rare fluctuation that brings it across the activation barrier ^1,5,10^. This picture highlights two points: 1) the long waiting period is to prepare the system for the transition period, 2) the transition period is caused by a rare fluctuation. To understand the physical mechanism of a rare event, we need to answer a few critical questions. 1) What is this rare fluctuation? 2) How does it cause transition? 3) What is the physical feature that distinguishes the dynamics during the transition and the waiting periods? This is challenging because we have little knowledge about this rare fluctuation except that it is the cause of the transition process—the only reliable clue is causality.

In this paper, we developed a rigorous procedure to identify the transition period of a rare event and parameterize its progression. This enabled us to carry out rigorous energy flow analysis of the transition period dynamics and understand its physical mechanism. Our results show that the rare fluctuation that initiates the transition period is a special phase-space condition (i.e. a point in the phase space) that triggers systematic energy flows in the system. The dynamics of the transition period are fundamentally different from the equilibrium fluctuations of the waiting period in that it ensures systematic energy flow from the non-reaction coordinates into the reaction coordinates, the few coordinates that fully decide the progress of a rare event. For a prototype of biomolecular isomerization reaction, the *C*_7*eq*_ → *C*_7*ax*_ transition of an alanine dipeptide in vacuum, the systematic energy flow during the transition period is achieved through direct transfer of kinetic energy from one coordinate to another by actions of inertial forces alone. This mechanism has not been discussed before and could represent a general mechanism for biomolecular conformational changes.

## Results

### Energy flow theory

Our main theoretical tool for analyzing the dynamics of a rare event is the energy flow theory we developed ^17,18^. This theory defines both potential (**PEFs**) and kinetic (**KEFs**) energy flows. The PEF through a coordinate *q*_*i*_ is its work ^18^:

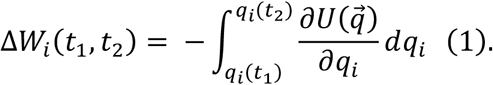

According to Eq. (1), Δ*W*_*i*_(*t*_1_, *t*_2_) is the change in the potential energy of the system due to the motion of *q*_*i*_ alone along a dynamic trajectory in the time interval [*t*_1_, *t*_2_]. It is a projection of the change in the total potential energy onto the motion of *q*_*i*_ and a measure of the cost of the motion of *q*_*i*_ in terms of potential energy. The change in the total potential energy of the system can be decomposed into PEFs through different coordinates: 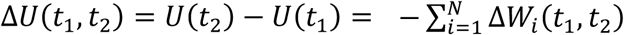, where the summation is over all the coordinates in the system. Along a similar line, the force from one coordinate *q*_*i*_ to another coordinate *q*_*j*_ is defined as: 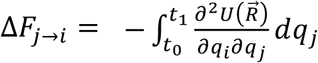, which measures how much of the force on *q*_*i*_ is due to the motion of *q*_*j*_ and its interaction with *q*_*i*_ ^18^.

The KEF through a coordinate *q*_*i*_ is ^17^:

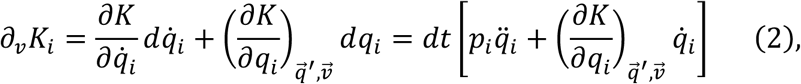

where *K* is the system kinetic energy, 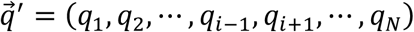 is the position vector in internal coordinates excluding *q*_*i*_, and 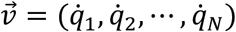 is the velocity vector. Since *∂*_*v*_*K*_*i*_ is the change in the system kinetic energy caused by changes in 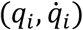 alone, which completely specifies the motion of *q*_*i*_, it rigorously defines the KEF through *q*_*i*_. It is equivalent to the change in the kinetic energy of *q*_*i*_ because it is always and only caused by motions of *q*_*i*_, even though we cannot define kinetic energy per coordinate in generalized coordinates because 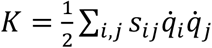, where *s*_*ij*_ is the coupling factor between 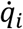 and 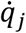 and a function of the structure of the system. Similar to PEFs, we can decompose Δ*K* as 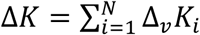 ^17^.

To gain mechanistic insights, we need to look at how the PEFs and KEFs of individual coordinates change with the progress of a rare event. We first project the PEF or KEF onto a projector *ξ*(Γ) that parameterizes the progress of a rare event, then average over the ensemble of reactive trajectories:

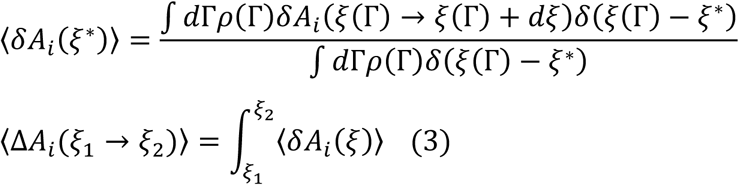

Here, *ρ*(Γ)*d*Γ is the probability of finding the system in an infinitesimal volume *d*Γ around a point Γ in phase space in the reactive trajectory ensemble; *δ*(*x*) is the Dirac δ-function; *δA*_*i*_ (*ξ*(Γ) → *ξ*(Γ) + *dξ*) is the change in *A*_*i*_ in a differential interval [*ξ*(Γ), *ξ*(Γ) + *dξ*) ; ⟨Δ*A*_*i*_ (*ξ*_1_ → *ξ*_2_)⟩ is the change in *A*_*i*_ in a finite interval [*ξ*_1_, *ξ*_2_], δ*A*_*i*_ can be either d*W*_*i*_ or *∂*_*v*_*K*_*i*_ ^17,18^. The ensemble of reactive trajectories consists of trajectories that cover the transition period but exclude the waiting period.

### Model System and Simulation Method

The process we study is the *C*_7*eq*_ → *C*_7*ax*_ isomerization of an alanine dipeptide in vacuum. This process is a prototype of conformational dynamics of proteins because alanine dipeptide is the smallest example of complex molecules. Here, we define a complex molecule as molecule whose non-reaction coordinates form a large enough heat bath for powering the reaction coordinates (**RC**s) to cross the activation barrier ^1,18-22^. In contrasts, small molecules need an external energy source for activation, such as buffer gas in gas-phase and solvents in solution-phase reactions. For this reason, we refer to the non-RCs in a complex molecule as the intra-molecular bath. The isomerization of an alanine dipeptide in vacuum carries some fundamental features that are unique to rare events in biomolecules but are absent in small molecules.

This isomerization process is mainly a rotation around the *ϕ* dihedral (Fig. 1). In previous studies, two backbone dihedrals *ϕ* and *θ*_1_ were identified as the RCs for this process, as they are sufficient for determining the committor value for any configuration of the system ^21,23^. Here, committor 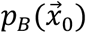 is the probability that a trajectory launched from the system configuration 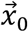, with momenta randomly drawn from equilibrium distribution, to reach the product basin before it visits the reactant basin ^1,19,24-28^. By definition, 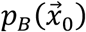 is the reaction probability of 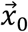 and rigorously parameterizes the progression of a rare event in the region of the configuration space that corresponds to *p*_*B*_ ∈ (0, 1).

**Figure 1:**
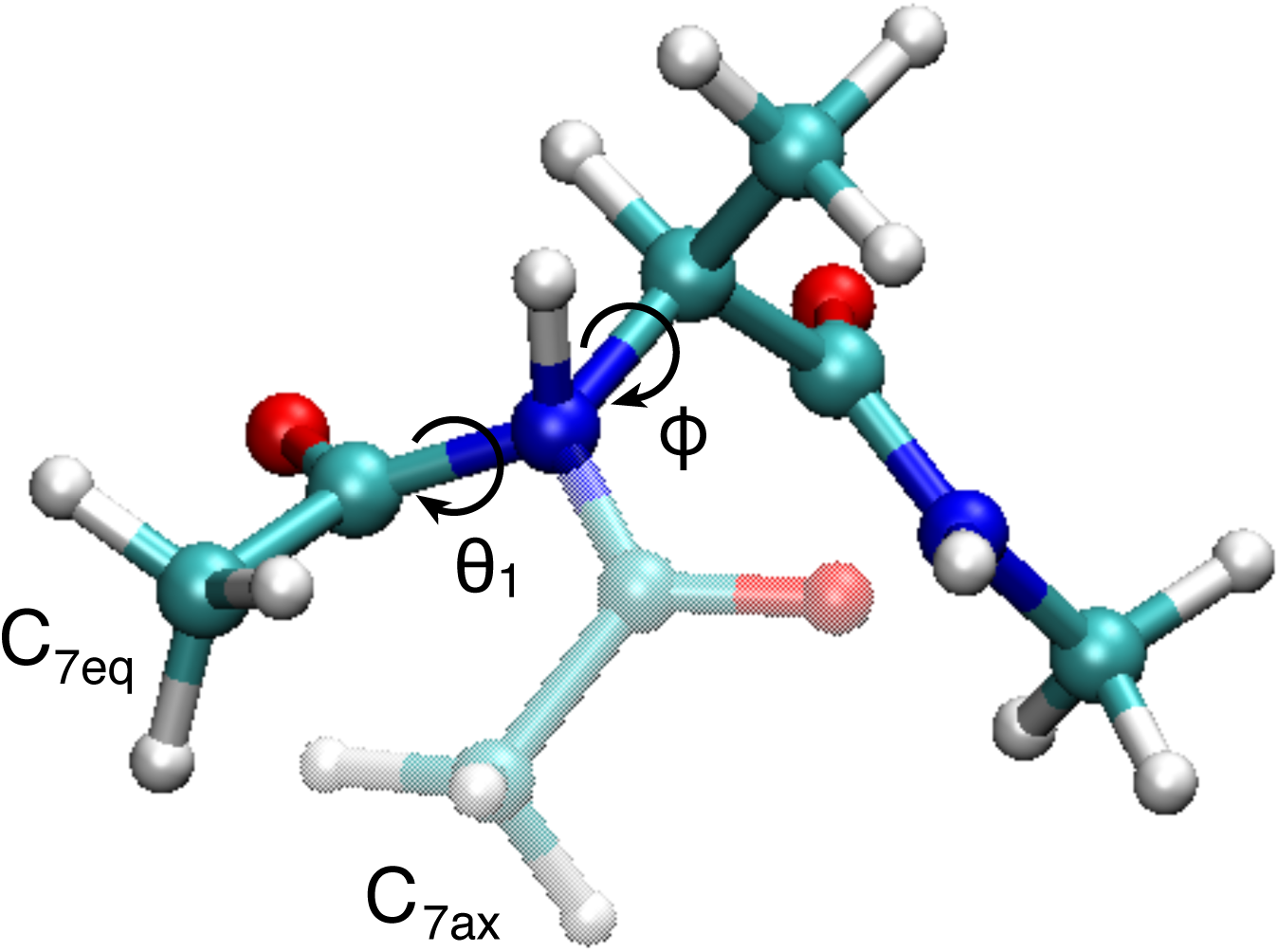
Two representative structures of an alanine dipeptide for the *C*_7*eq*_ (solid color) and *C*_7*ax*_ (semi-transparent color) states. The parts of the molecule that does not change in the *C*_7*eq*_ → *C*_7*ax*_ transition are completely overlapped between the two representations. The reaction coordinates *ϕ* and *θ*_1_ are labeled.

In previous energy flow analyses of this isomerization process ^17,18^, we found that *ϕ* needs to cross a potential energy barrier of ∼10 kJ⋅mol^-1^ in the region *p*_*B*_ ∈ (0, 1) and the energy required for crossing this barrier is mostly provided by KEF through *ϕ*. In contrast, *θ*_1_ receives ∼3 kJ⋅mol^-1^ of energy from other coordinates via PEF and uses this energy to help *ϕ* by directly transferring kinetic energy to *ϕ*. An intriguing question is: Where does the kinetic energy consumed by *ϕ* come from?

### An Example of the Rare Fluctuation That Causes Barrier Crossing

Figure 2 (upper panel) shows the time evolution of *ϕ* and *θ*_1_ along a reactive trajectory that contains both a failed and a successful barrier crossing. At *t* = 0.4 ps, *ϕ* reached a critical value: *ϕ* = 0.55. This value matches the value of *ϕ* at t = 1.03 ps when the successful barrier crossing starts, marked by committor beginning to rise above 0. Instead of moving forward and crossing the activation barrier, however, *ϕ* reversed its direction and went back to the reactant basin. After a brief stay there, *ϕ* moved towards the activation barrier again and reached the critical value of 0.55 at t = 1.03 ps. This time, it crossed the barrier successfully. Why did *ϕ* fail the first time but succeed the second time?

**Figure 2:**
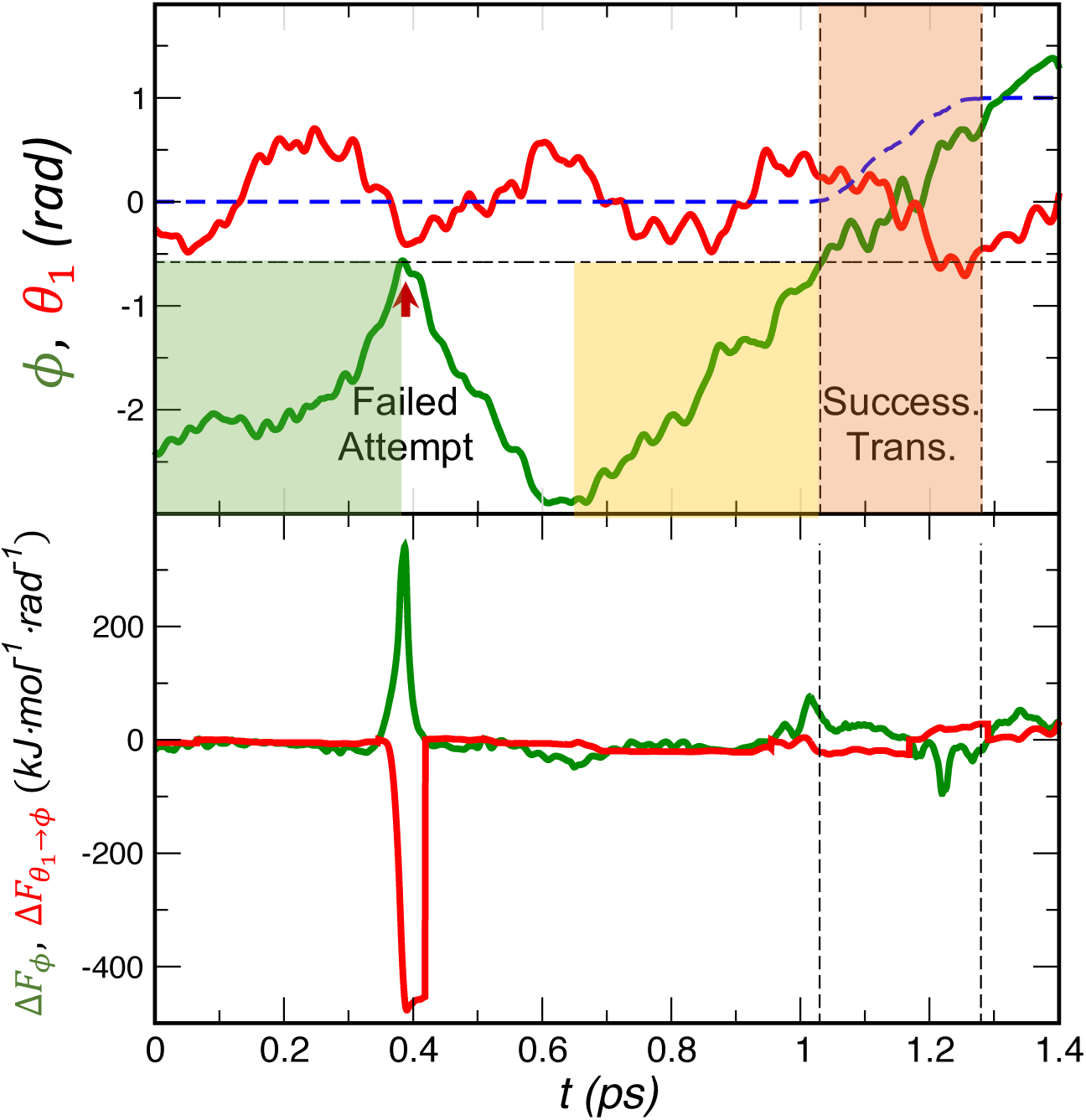
(**Upper**) Time evolution of *ϕ* (Green) and *θ*_1_ (Red) along a reactive trajectory that includes a failed attempt of barrier crossing marked by an arrow. Blue dashed line: time evolution of *p*_*B*_. The horizontal dashed line shows the critical value of *ϕ* that marks the onset of barrier crossing. The two vertical dashed lines mark the region of the successful barrier crossing. The two shaded rectangles mark the corresponding periods that precede *ϕ* reaching the critical value in the failed (Green) and successful (Yellow) barrier crossing respectively. The Brown shade mark the period for the successful barrier crossing. (**Lower**) Time evolution of the total force acting on *ϕ* (Δ*F*_*ϕ*_ ; Green) and the force from *θ*_1_ to *ϕ* (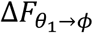; Red). Note the huge difference in the magnitudes of Δ*F*_*ϕ*_ and 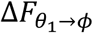 between the failed attempt around t = 0.4 ps and the successful transition during *t* ∈ [1, 1.3] ps.

Apparently, *ϕ* = 0.55 is a critical point for deciding the outcome of a barrier crossing. Inspecting the motion of *ϕ* gives the impression this is a stochastic and instantaneous decision because the motions of *ϕ* during the failed and successful barrier crossings do not show meaningful difference until *ϕ* = 0.55. Examination of quantities with more mechanistic significance, however, reveals that this decision is deterministic and it was made long before *ϕ* reaches 0.55.

Figure 2 (lower panel) shows the time evolution of 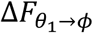, the force from *θ*_1_ to *ϕ*, along the same trajectory. In the failed attempt, 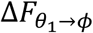 rises rapidly as *ϕ* approaches 0.55 and reaches a very high value at *ϕ* = 0.55. This force pushed *ϕ* away from the activation barrier, even though the other coordinates collectively applied a very large force (*F*_*ϕ*_) to push *ϕ* forward. In the successful attempt, in contrast, 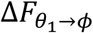 increases slowly, making its value at *ϕ* = 0.55 orders of magnitude smaller. Despite that *F*_*ϕ*_ is small this time, it is sufficient to push *ϕ* forward to cross the activation barrier. Based on these observations, the physical factor that makes the instantaneous decision on the outcome of a barrier crossing is the value of 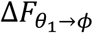 at *ϕ* = 0.55.

The value of 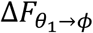 at *ϕ* = 0.55 is not instantaneously decided. Instead, it is determined by its evolution before *ϕ* reaches 0.55. Thus, the real cause of barrier crossing is the factor that determines this evolution process. Since Hamiltonian dynamics is deterministic, this evolution process is fully determined by the phase-space condition of its starting point. This specific starting point, therefore, decides the fate of an attempt at barrier crossing. The question is: Where is this specific starting point?

Figure 2 shows that, as *ϕ* approaches 0.55 during the second attempt, a systematic change in 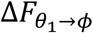 starts around t = 0.9 ps when *ϕ* is still well within the reactant basin. It is tempting to deduce that this is the starting point that we look for. However, this reasoning is not warranted because it is based on an empirical observation without justifications from a physically more rigorous ground. It is safe, however, to infer that a period of systematic dynamics starts somewhere between the time when *ϕ* returned to the reactant basin and the time it reached 0.55 again.

In the example above, barrier crossing is enabled by a proper value of 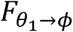 at *ϕ* = 0.55, which is determined by its evolution beforehand. This evolution appears systematic, pointing to a systematic component of the underlying dynamics that determines 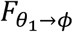. Since 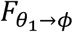 is unique to this example while dynamics is universal, the lesson is: the dynamics of the transition period has a systematic component. This systematic dynamics is distinct from equilibrium fluctuations that precede it because it serves a specific purpose: making the system ready at the onset of barrier crossing. In contrast, equilibrium fluctuations are thermal noise and do not serve any purpose. The key for successful barrier crossing is to maintain this systematic dynamics.

In Hamiltonian dynamics everything is fully determined by phase-space conditions, thus we need special phase-space conditions to maintain this systematic dynamics. Since this systematic process is determined by the phase-space point at its beginning, this point is the real cause of a successful barrier crossing. Reaching this special phase-space point is the rare fluctuation emphasized in the general picture of a rare event. The central questions are: 1) When does this systematic dynamics begin? 2) What does the system do during this period of systematic dynamics? To obtain rigorous answers, we need a comprehensive analysis of a rare event.

### Systematic Parameterization and Identification of the Transition Period

We consider a long Hamiltonian trajectory ***χ***(Γ(*t*), *t* ∈ [0, *T*]) that covers the entire course of one incidence of a rare event, from the moment when the system first enters the reactant basin at *t* = 0 until it finally leaves the reactant basin, crosses the activation barrier and reaches the product basin at time *t* = *T* (Fig. 3). Here, 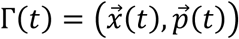 is the phase-space point on ***χ*** at time t; 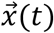 and 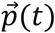 are the positions and momenta of all the degrees of freedom in the system respectively. The deterministic nature of Hamiltonian dynamics warrants that ***χ*** contains all the relevant information for understanding the mechanism of a rare event.

**Figure 3:**
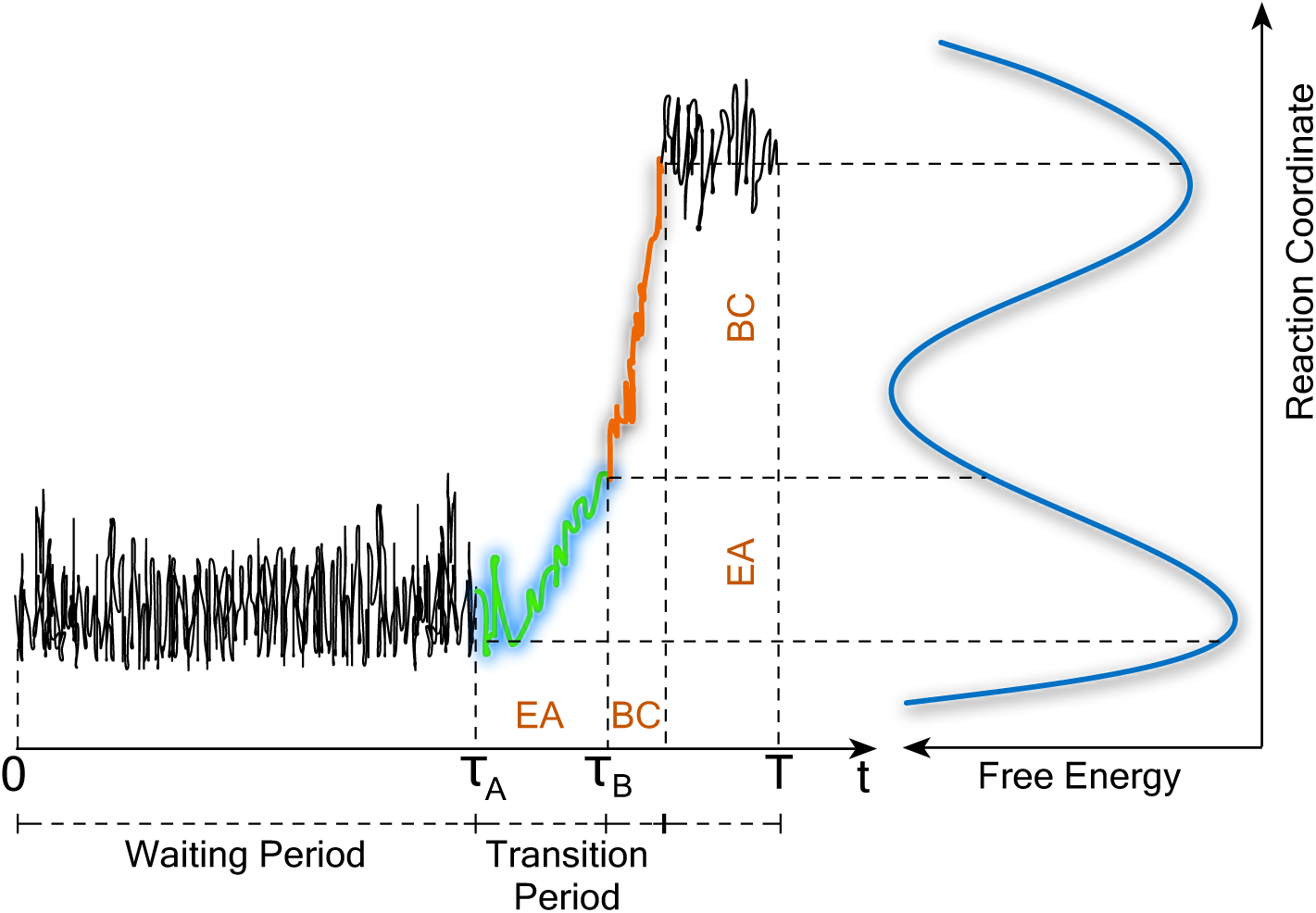
A schematic showing different phases of a dynamic trajectory that covers the entire course of one incidence of a rare event. The trajectory of the waiting period is shown as Black line, the trajectory of EA phase is shown in Green, and the trajectory of BC phase is shown in Orange. The schematic free energy profile along the reaction coordinate is shown in Blue. The vertical dashed lines mark the ranges of the three periods along the time axis; the horizontal dashed lines mark their ranges along the reaction coordinate axis.

This trajectory can be divided into two segments, the waiting period followed by the transition period. The waiting period is marked by equilibrium fluctuations. The transition period consists of two segments (Fig. 3): 1) the barrier crossing process that starts at *τ*_*B*_, 2) the systematic dynamics that precedes barrier crossing and ensures its success, which starts at *τ*_*A*_. The systematic dynamics in the time interval [*τ*_*A*_, *τ*_*B*_) is the essence of a rare event; it is enabled by the phase-space condition 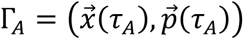 on ***χ*** at *τ*_*A*_. Our task is to identify *τ*_*A*_.

Committor can be computed independently, thus *τ*_*B*_ can be identified by finding the latest point on ***χ*** with 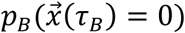 (i.e. among all the time t with 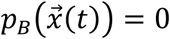, *τ*_*B*_ is the largest). Then *τ*_*A*_ can be located by tracing back in time from *τ*_*B*_ until we hit the point that is the necessary and sufficient condition for maintaining the systematic dynamics. The reason is the following.

A phase-space point Γ_*t*_(*t* ∈ [*τ*_*A*_, *τ*_*B*_]) on ***χ*** during [*τ*_*A*_, *τ*_*B*_] is a part of the systematic dynamics and determines all the dynamics after *t* and ensures it remains systematic. This makes Γ_*t*_ a sufficient condition for maintaining the systematic dynamics. In contrast, a point earlier than *τ*_*A*_ is not a sufficient condition because the systematic dynamics has not started yet--it cannot be maintained before it starts. The point at *τ*_*A*_ is therefore the first sufficient condition for maintaining the systematic dynamics. It is also the necessary condition because it is needed to start the systematic dynamics. Together, these make the point at *τ*_*A*_ (i.e. Γ_*A*_) the necessary and sufficient condition for maintaining the systematic dynamics.

In contrast, a point on ***χ*** after *τ*_*A*_ is a sufficient but not necessary condition. When we trace backward in time along ***χ*** from *τ*_*B*_, all the points on the way are sufficient conditions, but only the point at *τ*_*A*_ is also a necessary condition. Thus, we can identify *τ*_*A*_ as the first necessary condition we encounter as we trace back in time from *τ*_*B*_. The key for locating *τ*_*A*_, therefore, is a rigorous way to recognize the necessary condition, which requires a rigorous way to quantify the necessity of a condition for maintaining systematic dynamics.

The definition for statement A as a necessary condition for statement B is: if A is false, then B is false. If we want to show that a phase-space condition Γ_0_ is necessary for maintaining systematic dynamics, we need to show that if Γ_0_ is modified, systematic dynamics will be disrupted. Below is a procedure for testing necessity based on this idea.

For a phase-space condition 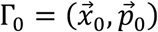, a small perturbation 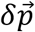 is added to its momenta to obtain a new condition 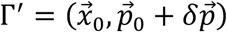. If the trajectory starting from Γ′ does not have a successful barrier crossing, it means that Γ′ cannot preserve systematic dynamics because systematic dynamics is defined as being able to ensure barrier crossing. This would mean Γ_0_ is a necessary condition because modifying it disrupts systematic dynamics. If the trajectory from Γ′ has a successful barrier crossing, it means that Γ′ can preserve systematic dynamics and Γ_0_ is not a necessary condition because Γ_0_ contains unnecessary components of at least the size of 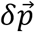.

Based on this reasoning, 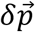 provides a measure of the level of necessity, but there is remaining uncertainty because 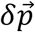 is not unique. To deal with this uncertainty, we use a set of perturbations 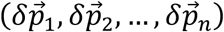 to generate an ensemble of perturbed conditions *E*(Γ_0_) = (Γ_1_, Γ_2_, …, Γ_*n*_). From each condition, we have a trajectory that either does or does not cross the barrier successfully. For the entire ensemble, we have a fraction of trajectories that have successful barrier crossings, which defines a probability *p*_*C*_(Γ_0_) that a perturbation to Γ_0_ will sustain systematic dynamics. We call *p*_*C*_(Γ_0_) the reaction capacity because it indicates the capability of Γ_0_ to launch reactive trajectories. Reaction capacity *p*_*C*_(Γ_0_) provides a statistical measure of how necessary Γ_0_ is for maintaining systematic dynamics; a larger value of *p*_*C*_(Γ_0_) means Γ_0_ is less necessary. Consequently, *p*_*C*_(Γ_0_) ≃ 0 means Γ_0_ is a necessary condition for maintaining systematic dynamics. Here, *p*_*C*_(Γ_0_) ≃ 0 means *p*_*C*_(Γ_0_) is infinitely close but not equal to 0.

Consider the phase-space condition at 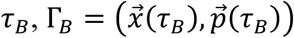, we expect *p*_*C*_(Γ_*B*_) ≃ 1. This is because 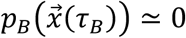, which means a complete change of 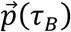 is required to disrupt the systematic nature of the dynamics. If we only perturb the momenta of Γ_*B*_ by a small amount, as required by the definition of *p*_*C*_, we expect most trajectories from the perturbed conditions should have successful barrier crossing, thus *p*_*C*_ (Γ_*B*_) ≃ 1. As we move backward in time, we expect that the *p*_*C*_ value of points on the trajectory on average will decrease--the necessity of the phase-space conditions along the trajectory increases. The first point with *p*_*C*_ ≃ 0 is the starting point of the transition period.

In the definition of *p*_*C*_, the magnitude of 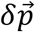 is the resolution for measuring the necessity for maintaining systematic dynamics; a larger magnitude of 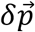 means lower resolution. It is an adjustable parameter that needs to be chosen empirically. For the current system, 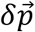 is generated by randomly sampling a value in the interval 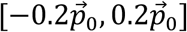 (i.e. *δ*p_*i*_ ∈ [−0.2p_0,*i*_, 0.2p_0,*i*_], *δ*p_*i*_ is the perturbation added to the momentum of coordinate *x*_*i*_) according to the uniform distribution. From each 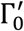 we launch a trajectory and check if it is reactive. In this way, we obtain the probability that trajectories launched from *E*(Γ_0_) are reactive, which is the value of *p*_*C*_ (Γ_0_).

### EA Precedes BC and Has Much Longer Duration

Figure 4 (left panel) shows the values of *p*_*C*_ and *p*_B_ along a typical reactive trajectory. They parameterize two consecutive segments; the segment with *p*_*C*_ ∈ (0, 1] precedes the segment with *p*_B_ ∈ (0, 1]; the portion with *p*_*C*_ ≃ 1 and the portion with *p*_B_ ≃ 0 coincide with each other. We call the first segment the energy activation (**EA**) phase and the second segment the barrier crossing (**BC**) phase (Fig. 3). The combination of EA and BC forms the transition period. The EA phase lasts more than 1 ps and *p*_*C*_ experiences several turns of back-and-forth changes in its course of increasing from 0 to 1. In contrast, BC lasts less than 200 fs and *p*_*B*_ increases monotonically ^29^. Figure 4 (right panel) shows that on average the duration of EA is about 5 times longer than the duration of BC, suggesting that EA is more complex and more important for the *C*_7*eq*_ → *C*_7*ax*_ transition. Both distributions can be fit to a log-normal form.

**Figure 4:**
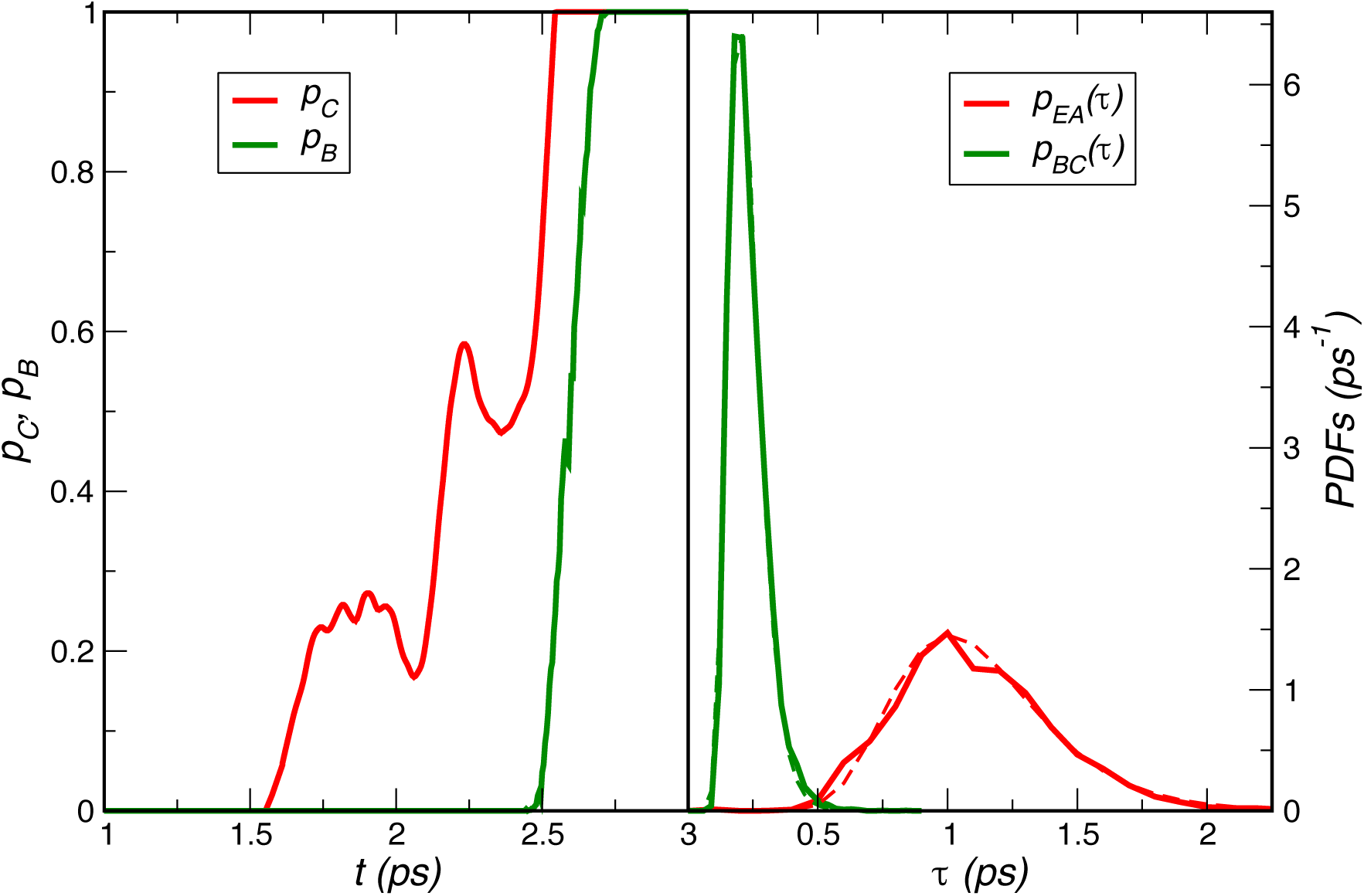
(***Left***) The values of *p*_*C*_ (Red) and *p*_*B*_ (Green) along a typical reactive trajectory. (***Right***) The probability density functions for the durations of the EA (*τ*_*EA*_) and BC (*τ*_BC_) phases. The PDFs are generated from 12,000 reactive trajectories. The duration of EA is much longer than that of BC. The dashed lines are the log-normal fits: 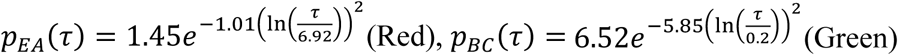.

### EA is Dominated by Direct Transfer of Kinetic Energy

Figure 5 shows the PEFs and KEFs through all the coordinates in the system during both EA and BC. To calculate energy flows during EA, we used *p*_*C*_ as the projector. For the BC phase, the projector is *p*_*B*_. The energy flows for the BC phase were shifted so that the energy flows at *p*_*C*_ = 1 and p_*B*_ = 0 are continuous because these two regions coincide with each other.

**Figure 5:**
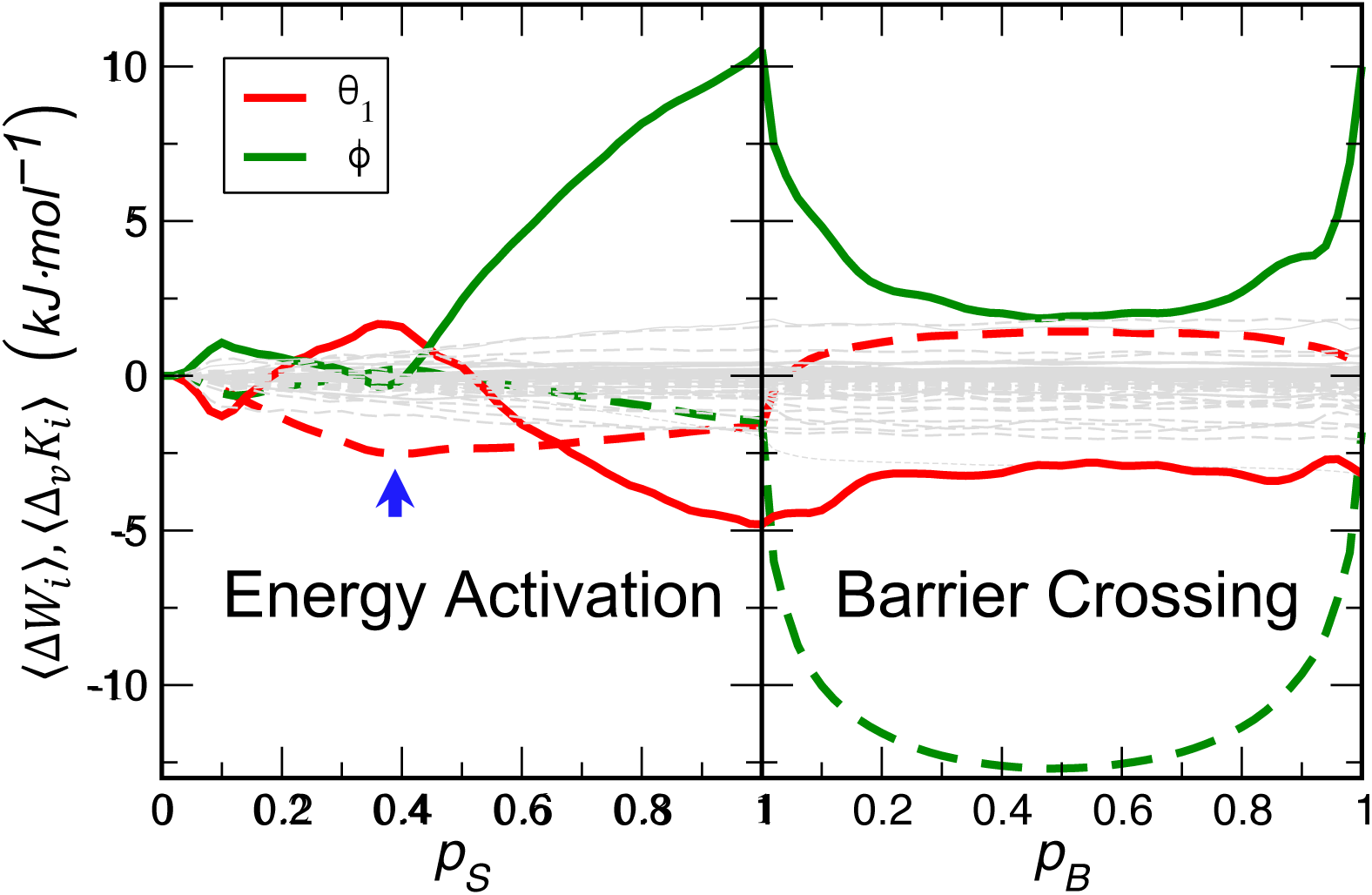
The PEFs (⟨Δ*W*_*i*_⟩ *i* = *ϕ, θ*_1_; dashed lines) and KEFs (⟨Δ_*v*_*K*_*i*_⟩ *i* = *ϕ, θ*_1_; solid lines) during the EA phase (left panel) and the BC phase (right panel). The Blue arrow marks the top of the barrier on the path of *θ*_1_ during energy activation. The gray dashed lines are the PEFs and KEFs of the other coordinates (58 in total) in the system.

As expected, only the two dominant reaction coordinates, *ϕ* and *θ*_1_, experience significant energy flows during EA. The KEFs of *ϕ* and *θ*_1_ are high and of opposite sign--*ϕ* gains while *θ*_1_ loses kinetic energy. The dependence of ⟨Δ_*v*_*K*_*ϕ*_⟩ and 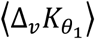 on *p*_*C*_ are like mirror images of each other except that the magnitude of the latter is about 50% of the former. Since the PEFs through both coordinates are much lower than the KEFs, this means the kinetic energy attained by *ϕ* is mainly through direct transfer of kinetic energy into *ϕ*,∼50% from *θ*_1_ and the other ∼50% from the intra-molecular bath. This process is realized via the actions of inertial forces instead of forces derived from potential energy according to the general mechanism of KEFs discovered in ref. ^17^ Throughout EA, kinetic energy accumulates in *ϕ* and reaches ∼10 kJ/mol at *p*_*C*_ = 1, about the same amount as the energy *ϕ* consumed while crossing the activation barrier. Therefore, the kinetic energy consumed during BC, first identified in ref. ^18^, is supplied by the accumulation during EA. To summarize, the main feature of the dynamics of EA phase is that kinetic energy systematically flows into and accumulates in the dominant RC, a process mainly administered by inertial forces. To our knowledge, this mechanism has not been discussed before and is unique to biomolecules.

Another essential feature of EA is the timing of different energy flows. If we consider the starting point of a systematic energy flow as when it rises above the noise level defined by the energy flows of all non-RCs (the cloud of gray lines in Fig. 5), then among all the systematic energy flows during EA, the PEF of *θ*_1_ starts the earliest. While the PEF of *ϕ* is negligible, *θ*_1_ needs to cross a moderate barrier of ∼2.5 kJ/mol at *p*_*C*_ ≃ 0.4. This is also the starting point of the systematic KEFs of *ϕ* and *θ*_1_, with the nearly perfect correlation between them pointing to a tight synergy between the motions of *ϕ* and *θ*_1_. Since proper KEF into *ϕ* is the necessary condition for BC, this means it is impossible for *ϕ* to start BC before *θ*_1_ crossed its own barrier and became ‘ready’ to help *ϕ* ^17^. Therefore, *θ*_1_ acts as a gating mechanism on *ϕ*. This effect also shows up in the trajectory: *θ*_1_ was in a wrong position during the failed attempt at t = 0.4 ps in Fig. 2 (upper panel), which is the reason that *ϕ* is pushed back by 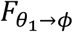 (Fig. 2). This precise coordination between *ϕ* and *θ*_1_ highlights the intricate mechanism of EA.

### A Physical Picture of Rare Events in Biomolecules

The results above suggest an intriguing physical picture of a rare event in a complex biomolecule that obeys Hamiltonian dynamics. A rare event has three phases: the waiting period, the EA phase and the BC phase. Each phase has distinctive features. During the waiting period, the system searches randomly in the phase space until it encounters a special phase-space condition that can initiate systematic energy flows. This is the moment that waiting ends and EA starts. During EA, energy flows from the intra-molecular bath into the RCs and accumulates there. The progress of EA is gauged by the amount of energy accumulated in RCs. When energy accumulation in RCs reaches its maximum, EA ends and BC starts. In this phase, the energy accumulated in RCs during EA is consumed to power barrier crossing.

Formulation of this picture was enabled by introducing *p*_*C*_(Γ_0_) as a measure of how necessary Γ_0_ is for maintaining systematic dynamics and energy flow, a rather abstract definition motivated by logical deduction. But *p*_*C*_(Γ_0_) also has an intuitive meaning based on the concept of reactive trajectory. Since *p*_*C*_(Γ_0_) is the probability that a trajectory launched from 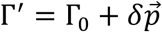 is reactive, it is the probability that a reactive trajectory passes through the close neighborhood around Γ_0_, an “ellipsoid” of “radius” 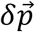 centered at Γ_0_. Therefore, *p*_*C*_(Γ_0_) is a measure of the density of reactive trajectories in the close neighborhood around Γ_0_. A small *p*_*C*_(Γ_0_) indicates a low density and a large *p*_*C*_(Γ_0_) indicates a high density. Since 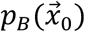 is essentially 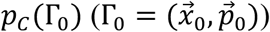 with 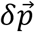 as large as 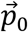, even small value of *p*_*B*_ indicates high density of reactive trajectories. This is why the regions of *p*_*C*_ ≃ 1 and p_B_≃ 0 coincide at the boundary between EA and BC (Fig. 4, left panel).

The alternative meaning of *p*_*C*_(Γ_0_) above leads to a physical picture of a rare event from a different perspective. Throughout the entire waiting period *p*_*C*_ stays 0, thus the system is searching in non-reactive regions of the phase space—regions with no reactive trajectories. This search is random because the dynamics during the waiting period is thermal fluctuations. This random search stops when the system encounters a phase-space condition Γ_0_ with p_*C*_(Γ_0_) ≃ 0 by pure chance. These phase-space conditions mark the region with low-density of reactive trajectories, thus they are the starting points of reactive trajectories and the entrance points to the reactive region—the phase-space region occupied by reactive trajectories. They are rare and scattered through the vast non-reactive region, thus the chance for the system to come across them is small, making the search and waiting long. In the EA phase, *p*_*C*_ increases from 0 to 1, thus the system moves from region with low-density to region with high density of reactive trajectories. This increase in reactive trajectory density continues into the BC phase and peaks at *p*_*B*_ = 0.5, i.e. the transition state region, confirming its status as the dynamic bottleneck ^1^. Afterwards, the reactive trajectory density starts to decrease gradually as *p*_*B*_ passes 0.5 and approaches 1.

## Discussions

In this paper, we developed a new metric, the reaction capability *p*_*C*_, that parameterizes the progress of the EA phase of a rare event. Together with committor *p*_B_ that parameterizes the BC phase, they provide rigorous parameterization of the entire transition period and enable us to identify its starting point and carry out rigorous analysis of its dynamics. Our results showed that EA precedes BC and has much longer duration; the energy accumulation during EA is a necessary condition for BC and critical for the function of biomolecules. Therefore, understanding the mechanism of EA is critical for understanding the mechanism of rare events in biomolecules.

Surprisingly, in contrast to the BC phase dominated by PEFs, the EA phase is dominated by kinetic energy that systematically flows from the intra-molecular bath to the RC and accumulates there. From our analysis on the relation between kinetic and potential energy flows in ref. ^17^, this fact means that the systematic KEFs during EA are achieved through direct transfer of kinetic energy from one coordinate to another under the actions of inertial forces rather than forces derived from potential energy. This is made possible by the special structural features of biomolecules that introduce structural couplings between different coordinates that synergize with the activation process to sustain systematic transfer and accumulation of kinetic energy. This may be the key to the unique functions of biomolecules that are not possible in small molecules. Although large proteins may involve more sophisticated mechanisms for EA, systematic transfer and accumulation of kinetic energy is likely an important component.

This effect cannot be explained by transition state theory or Kramers theory: the former skips dynamics and the latter is based on an equation of motion, the Langevin equation, that does not support energy transfer administered by the inertial forces ^2,10,11^. This might be the reason that application of Kramers theory to understanding observations from single molecule spectroscopy has met significant challenges. For example, Neupane et al found that the barrier height and diffusion constant extracted from single molecule experiments on protein folding are inconsistent with the measured transition path time ^4,13,30-32^. Antoniou and Schwartz found that velocities of some key residues play a critical role in determining the progress of the hydride transfer reaction in lactate dehydrogenase, suggesting the importance of momentum space ^33^. These observations cast doubt on whether protein conformational dynamics and folding can be adequately described by Kramers theory in the spatial diffusion regime ^4,11,13^. To our knowledge, this systematic transfer and accumulation of kinetic energy steered by inertial forces has not been discussed in the well-studied process of intra-molecular vibrational redistribution either ^34-39^.

The picture discussed here is based on Hamiltonian dynamics, which is always valid because any system will obey Hamiltonian dynamics if all of its degrees of freedom are explicitly considered (e.g. both the buffer gas and the reactants in a gas phase reaction are explicitly included in the system Hamiltonian). Therefore, this picture is generally valid as long as quantum effects can be neglected. Although the example we examined is in vacuum whereas proteins work in solutions, the picture and approach presented here provide a foundation for understanding activation of biomolecules in solutions. This is because we developed a general and rigorous approach for understanding the intra-molecular bath, which is an extra layer of functional machinery that plays a critical role in the activation dynamics and functions of biomolecules. The intra-molecular bath is the reason that a biomolecule can organize systematic energy flows of much higher magnitude than what is possible in a small molecule, which is key for achieving the unique functions of a protein (e.g. enzymatic catalysis) that are not possible for small molecules. Solvent molecules will have strong and extensive interactions with the intra-molecular bath, whereas their interactions with the protein RCs could be more limited due to the small number of RCs. On the other hand, our previous study showed that solvents play a dominant role of an isomerization of an alanine dipeptide in water, but that could be due to the small size of the solute so that the RC is fully exposed to the solvents ^21^. It remains to be seen whether solvents should be treated on an equal footing as the protein using full Hamiltonian representation, or they should be treated in a statistical manner as an additional bath for providing energy input into the intra-molecular bath to augment the energy flow of the latter.

Finally, we note that the reaction capacity *p*_*C*_(Γ_0_) defined here is fundamentally different from the extended committor by Antoniou and Schwartz ^33^, which was based on the stochastic separatrix idea derived from Kramers theory ^40,41^. Calculating the extended committor involves randomizing all the momenta except those conjugate to the RCs by sampling from a Boltzmann distribution.

## Methods

All simulations were performed using the molecular dynamics software suite GROMACS ^42^with transition path sampling implemented. Amber 94 force field was used to facilitate comparison with previous results ^17,18,23,43,44^. The structure of the alanine dipeptide was minimized using steepest descent algorithm and heated to 300 K using velocity rescaling with a coupling constant of 0.2 ps. The system was then equilibrated for 200 ps and no constraints were applied. The time step of integration was 1 fs. Basin *C*_7*eq*_ is defined as −180° < *ϕ* < −55° and 0° < *ψ* < 180°; basin *C*_7*ax*_ is defined as 50° < *ϕ* < 100° and −180° < *ψ* < 0°. We used transition path sampling method to generate the ensemble of reactive trajectories between these two basins that are used in all the analyses discussed here ^1^. All the averaged quantities discussed in the text were averaged over 12,000 trajectories.

## Notes

### Competing Interest Statement

The authors have declared no competing interest.

## References

1 Bolhuis, P. G., Chandler, D., Dellago, C. and Geissler, P. L. Transition path sampling: Throwing ropes over rough mountain passes, in the dark. Ann. Rev. Phys. Chem. 53, 291–318 (2002).

2 Garcia-Viloca, M., Gao, J., Karplus, M. & Truhlar, D. G. How enzymes work: Analysis by modern rate theory and computer simulations. Science 303, 186–195, doi:DOI 10.1126/science.1088172 (2004).

3 Schwartz, S. D. & Schramm, V. L. Enzymatic transition states and dynamic motion in barrier crossing. Nat Chem Biol 5, 551–558, doi:10.1038/nchembio.202 (2009).

4 Zhang, B. W., Jasnow, D. & Zuckerman, D. M. Transition-event durations in one-dimensional activated processes. J Chem Phys 126, 074504, doi:10.1063/1.2434966 (2007).

5 Pollak, E. & Talkner, P. Reaction rate theory: What it was, where is it today, and where is it going? Chaos 15, 026116, doi:Artn 02611610.1063/1.1858782 (2005).

6 Wigner, E. The transition state method. Trans. Farady Soc. 34, 29–41 (1938).

7 Grote, R. F. & Hynes, J. T. The Stable States Picture of Chemical-Reactions .2. Rate Constants for Condensed and Gas-Phase Reaction Models. J Chem Phys 73, 2715–2732, doi:Doi 10.1063/1.440485 (1980).

8 Hynes, J. T. & Grote, R. F. Models for Atom Transfer and Isomerization-Reactions in Liquids. Abstr Pap Am Chem S 180, 258–Phys (1980).

9 Berne, B. J., Borkovec, M. & Straub, J. E. Classical and Modern Methods in Reaction-Rate Theory. J Phys Chem-Us 92, 3711–3725, doi:DOI 10.1021/j100324a007 (1988).

10 Hanggi, P., Talkner, P. & Borkovec, M. Reaction-Rate Theory - 50 Years after Kramers. Rev Mod Phys 62, 251–341, doi:Doi 10.1103/Revmodphys.62.251 (1990).

11 Kramers, H. A. Brownian motion in a field of force and the diffusion model of chemical reactions. Physica VII, 284–304 (1940).

12 Pechukas, P. in Dynamics of Molecular Collisions Part B Vol. 2 (ed W. H. Miller) Ch. 6, 269 (Plenum, 1976).

13 Chaudhury, S. & Makarov, D. E. A harmonic transition state approximation for the duration of reactive events in complex molecular rearrangements. J Chem Phys 133, 034118, doi:10.1063/1.3459058 (2010).

14 Chung, H. S., Louis, J. M. & Eaton, W. A. Experimental determination of upper bound for transition path times in protein folding from single-molecule photon-by-photon trajectories. Proc Natl Acad Sci U S A 106, 11837–11844, doi:10.1073/pnas.0901178106 (2009).

15 Chung, H. S., McHale, K., Louis, J. M. & Eaton, W. A. Single-molecule fluorescence experiments determine protein folding transition path times. Science 335, 981–984, doi:10.1126/science.1215768 (2012).

16 Schramm, V. L. & Schwartz, S. D. Promoting Vibrations and the Function of Enzymes. Emerging Theoretical and Experimental Convergence. Biochemistry 57, 3299–3308, doi:10.1021/acs.biochem.8b00201 (2018).

17 Li, H. & Ma, A. Kinetic energy flows in activated dynamics of biomolecules. J. Chem. Phys. 153, 094109, doi:doi: 10.1063/5.0020275 (2020).

18 Li, W. & Ma, A. Reaction mechanism and reaction coordinates from the viewpoint of energy flow. J Chem Phys 144, 114103, doi:10.1063/1.4943581 (2016).

19 Du, R., Pande, V. S., Grosberg, A. Y., Tanaka, T. & Shakhnovich, E. S. On the transition coordinate for protein folding. J Chem Phys 108, 334–350, doi:Doi 10.1063/1.475393 (1998).

20 Li, W. & Ma, A. Recent developments in methods for identifying reaction coordinates. Mol Simul 40, 784–793, doi:10.1080/08927022.2014.907898 (2014).

21 Ma, A. & Dinner, A. R. Automatic method for identifying reaction coordinates in complex systems. J Phys Chem B 109, 6769–6779, doi:10.1021/jp045546c (2005).

22 Berezhkovskii, A. & Szabo, A. One-dimensional reaction coordinates for diffusive activated rate processes in many dimensions. J Chem Phys 122, doi:Artn 01450310.1063/1.1818091 (2005).

23 Bolhuis, P. G., Dellago, C. & Chandler, D. Reaction coordinates of biomolecular isomerization. P Natl Acad Sci USA 97, 5877–5882, doi:Doi 10.1073/Pnas.100127697 (2000).

24 Ryter, D. On the Eigenfunctions of the Fokker-Planck Operator and of Its Adjoint. Physica A 142, 103–121, doi:Doi 10.1016/0378-4371(87)90019-7 (1987).

25 Onsager, L. Initial recombination of ions. Phys. Rev. 54, 554–557 (1938).

26 Berezhkovskii, A. M. & Szabo, A. Diffusion along the splitting/commitment probability reaction coordinate. J Phys Chem B 117, 13115–13119, doi:10.1021/jp403043a (2013).

27 Tiwary, P. & Berne, B. J. Spectral gap optimization of order parameters for sampling complex molecular systems. P Natl Acad Sci USA 113, 2839–2844, doi:10.1073/pnas.1600917113 (2016).

28 Peters, B. & Trout, B. L. Obtaining reaction coordinates by likelihood maximization. J Chem Phys 125, 054108, doi:Artn 05410810.1063/1.2234477 (2006).

29 Li, W. J. & Ma, A. Reducing the cost of evaluating the committor by a fitting procedure. J. Chem. Phys. 143, 174103 (2015).

30 Neupane, K. et al. Direct observation of transition paths during the folding of proteins and nucleic acids. Science 352, 239–242, doi:10.1126/science.aad0637 (2016).

31 Manuel, A. P., Lambert, J. & Woodside, M. T. Reconstructing folding energy landscapes from splitting probability analysis of single-molecule trajectories. Proc Natl Acad Sci U S A 112, 7183–7188, doi:10.1073/pnas.1419490112 (2015).

32 Woodside, M. T. et al. Direct measurement of the full, sequence-dependent folding landscape of a nucleic acid. Science 314, 1001–1004, doi:10.1126/science.1133601 (2006).

33 Antoniou, D. & Schwartz, S. D. Phase Space Bottlenecks in Enzymatic Reactions. J Phys Chem B 120, 433–439, doi:10.1021/acs.jpcb.5b11157 (2016).

34 Jang, S. M., Zhao, M. S. & Rice, S. A. Comment on the Rate of Isomerization in Molecules with a Symmetrical Triple Well Potential. J Chem Phys 97, 8188–8196, doi:Doi 10.1063/1.463441 (1992).

35 Zhao, M. S. & Rice, S. A. Comment on the Classical-Theory of the Rate of Isomerization. J Chem Phys 97, 943–951, doi:Doi 10.1063/1.463197 (1992).

36 Buchenberg, S., Leitner, D. M. & Stock, G. Scaling Rules for Vibrational Energy Transport in Globular Proteins. J Phys Chem Lett 7, 25–30, doi:10.1021/acs.jpclett.5b02514 (2016).

37 Leitner, D. M. in Annu Rev Phys Chem Vol. 59 Annual Review of Physical Chemistry 233–259 (2008).

38 Fujisaki, H. & Straub, J. E. Vibrational energy relaxation in proteins. Proc Natl Acad Sci U S A 102, 6726–6731, doi:10.1073/pnas.0409083102 (2005).

39 Botan, V. et al. Energy transport in peptide helices. Proc Natl Acad Sci U S A 104, 12749–12754, doi:10.1073/pnas.0701762104 (2007).

40 Klosek, M. M., Matkowsky, B. J. & Schuss, Z. The Kramers Problem in the Turnover Regime: The Role of the Stochastic Separatrix. Marz 95, 331 –337 (1991).

41 Schuss, Z. & Spivak, A. The Exit Distribution on the Stochastic Separatrix in Kramers’ Exit Problem. SIAM J. App. Math. 62, 1698 –1711 (2002).

42 Hess, B., Kutzner, C., van der Spoel, D. & Lindahl, E. GROMACS 4: Algorithms for highly efficient, load-balanced, and scalable molecular simulation. J Chem Theory Comput 4, 435–447, doi:10.1021/ct700301q (2008).

43 Case, D. A. et al. Amber 5.0. University of California, San Francisco (1997).

44 Pearlman, D. A. et al. Amber, a Package of Computer-Programs for Applying Molecular Mechanics, Normal-Mode Analysis, Molecular-Dynamics and Free-Energy Calculations to Simulate the Structural and Energetic Properties of Molecules. Comput Phys Commun 91, 1–41, doi:Doi 10.1016/0010-4655(95)00041-D (1995).

